# Visual experience regulates the intrinsic excitability of visual cortical neurons to maintain sensory function

**DOI:** 10.1101/402768

**Authors:** Alexander P.Y. Brown, Lee Cossell, Troy W. Margrie

**Affiliations:** The Sainsbury Wellcome Centre for Neural Circuits and Behaviour, University College London, 25 Howland Street, London W1T 4JG, United Kingdom

## Abstract

This *in vivo* study shows that both intrinsic and sensory-evoked synaptic properties of layer 2/3 neurons in mouse visual cortex are modified by ongoing visual input. Following visual deprivation, intrinsic properties are significantly altered, although orientation selectivity across the population remains unchanged. We therefore suggest that cortical cells adjust their intrinsic excitability in an activity-dependent manner to compensate for changes in synaptic drive and maintain sensory network function.

---

Recordings from neurons *in vitro* show that the biophysical profile of a given cell can be influenced by recent activity^1-4^ and that such intrinsic plasticity can alter the response to input^1,5,6^. In the mammalian brain, while experience-dependent changes in cellular excitability are partly dependent on changes at synapses^7-9^, no study has yet investigated whether externally-driven activity might drive intrinsic changes *in vivo*, or how this could be used to optimize or maintain physiological function, for example a particular output, such as overall firing rate^10,11^.

To explore this question, we performed whole-cell recordings from layer 2/3 regular-spiking neurons in primary visual cortex of anaesthetized mice that, post-weaning, had either been housed under normal lighting conditions (control: 12h light/12h dark, n=150 cells) or in complete darkness for up to five weeks (visually deprived, V.D; n=40 cells). We observed that visually deprived mice had a significantly depolarized resting membrane potentials (median control: −76.8mV (interquartile range, IQR: 9.0mV) vs. V.D.: −71.6mV (IQR: 9.6mV); p=0.0019 Wilcoxon rank-sum test; **Fig. 1a,b**) and spike threshold (control: −27.9mV (IQR: 10.4mV) vs. V.D.: −24.3mV (IQR: 9.7mV); p=1.4×10^−4^; **Fig. 1a,c**) compared to control mice. Although the resting membrane potential and spike threshold both shifted to more depolarized potentials in the absence of visual input, the distance from rest to threshold did not change (control: 49.1mV (IQR: 11.7mV) vs. V.D.: 48.9mV (IQR: 15.4mV); p=0.64; **Fig. 1d**). Furthermore, neurons in visually deprived animals exhibited an increased input resistance to depolarizing (control: 87.0MΩ (IQR: 51.3MΩ), n=128 vs. V.D.: 104.7MΩ (IQR: 68.1MΩ), n=35; p=0.019; **Fig. 1e,f**) but not hyperpolarizing current injections (control: 51.4MΩ (IQR: 30.8MΩ), n=150 vs. V.D.: 52.9MΩ (IQR: 36.6MΩ), n=40; p=0.71). and subsequently, showed lower rheobase values (control: 275pA (IQR: 125pA) vs. V.D.: 225pA (IQR: 87.5pA); p=0.0086; **Fig. 1g**) suggesting that active conductances responsible for rectification in L2/3 cells are altered following changes in visual input.

**Figure 1.**
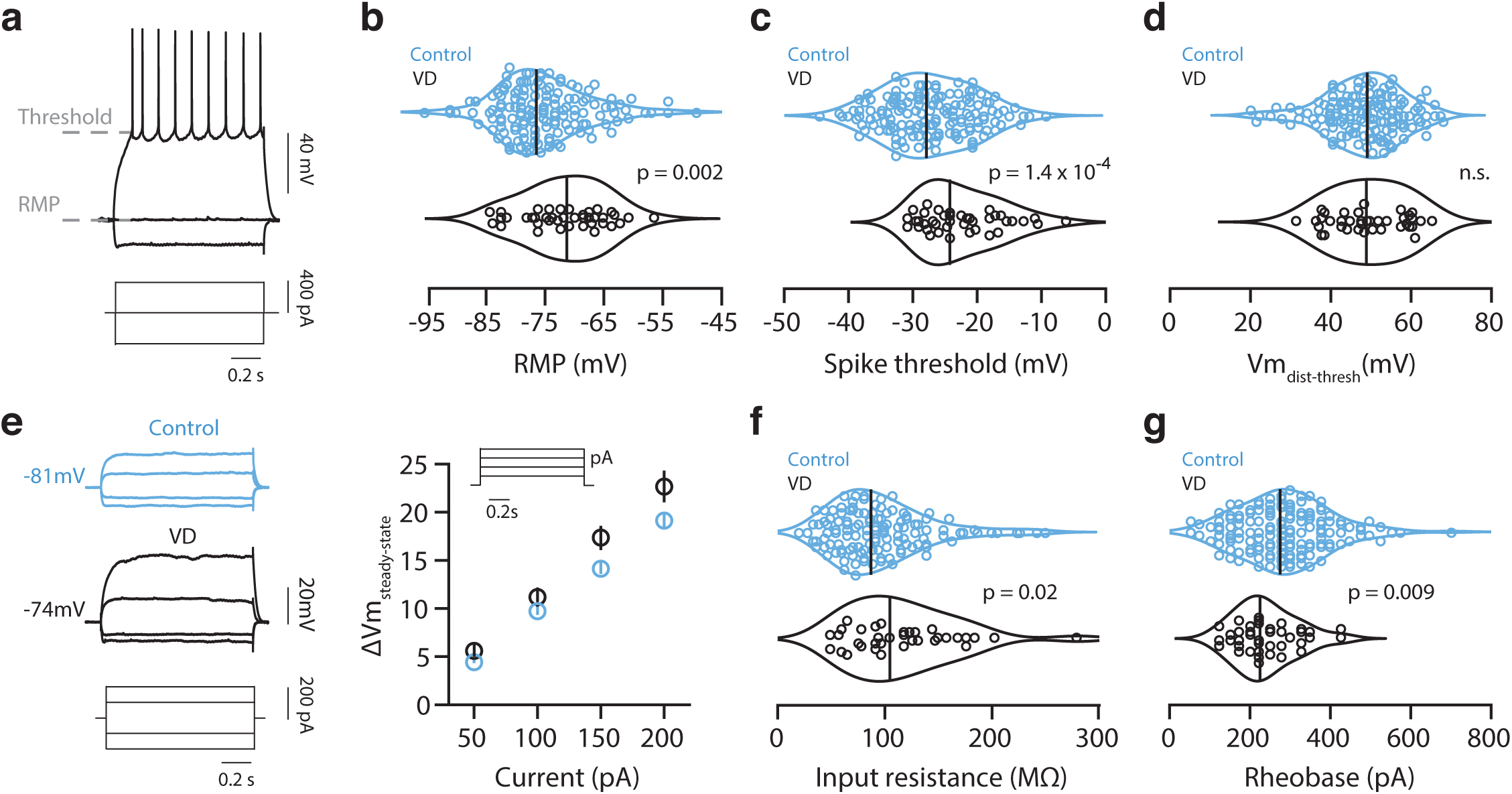
Intrinsic membrane properties of layer 2/3 pyramidal cells are altered in visually deprived mice. Example membrane potential traces evoked by hyperpolarizing and depolarizing current steps. Violin plots of resting membrane potential (**b**), spike threshold at rheobase (**c**), and distance between resting membrane potential and spike threshold (**d**) recorded in control (blue) and visually deprived (black) mice. (**e**) Left, example membrane potential traces in response to hyperpolarizing and depolarizing current steps recorded in control and visually deprived mice. Right, graph showing the amplitude of the steady state membrane potential evoked by depolarizing current steps. Median and S.E.M. shown. (**f**, **g**) Violin plots of input resistance (**f**) and rheobase (**g**). (**b**-**d**,**g** control n=150 vs. V.D. n=40 cells; **f**, control n=128 vs. V.D. n=35 cells). Vertical bars on the violin plots indicate median values.

To understand how such cellular changes might impact function, in a subset of cells we also quantified the visually-evoked synaptic and spiking responses to moving grating stimuli (control: n=128; V.D.: n=35 neurons) (**Fig. 2a**). For each grating direction, we quantified the average evoked synaptic depolarization (Vm_0_; **Fig. 2a,b**), the modulation of the synaptic response (Vm_1_; **Fig. 2a,c**) and the mean spike rate (**Fig. 2d**). Although there was little change in the overall amount of depolarization (Vm_0_ at preferred spiking direction: control 11.2mV (IQR: 6.8mV) vs. V.D. 10.1mV (IQR: 3.6mV), p=0.10; Wilcoxon rank-sum test; **Fig. 2b**), there was a significant decrease in the amount of membrane potential modulation (Vm_1_ at preferred, control 11.0mV (IQR: 7.9mV) vs. V.D. 7.2mV (IQR: 5.8mV), p=9.1×10^−4^; **Fig. 2c**). Cells recorded from animals with reduced visual input also show a reduction in evoked spike rates (spike rate at preferred, control 2.15Hz (IQR: 4.86Hz) vs V.D. 0.95Hz (IQR: 2.34Hz); p=0.0058; **Fig. 2d**), and the number of neurons without visually-evoked spikes significantly increased compared to control (control: 1/128 vs. V.D: 7/35; p < 0.0001, Fisher-exact test). Thus, reducing visual input alters not only intrinsic set points such as resting membrane potential and threshold but also modifies both the synaptic and spiking responsiveness of cells. It is not possible to discern from these data whether the reduced input modulation is due to a decrease in driving force caused by more deploarised potentials or synaptic strength. In any case, increased input resistance is expected to at least partly counteract these effects.

**Figure 2.**
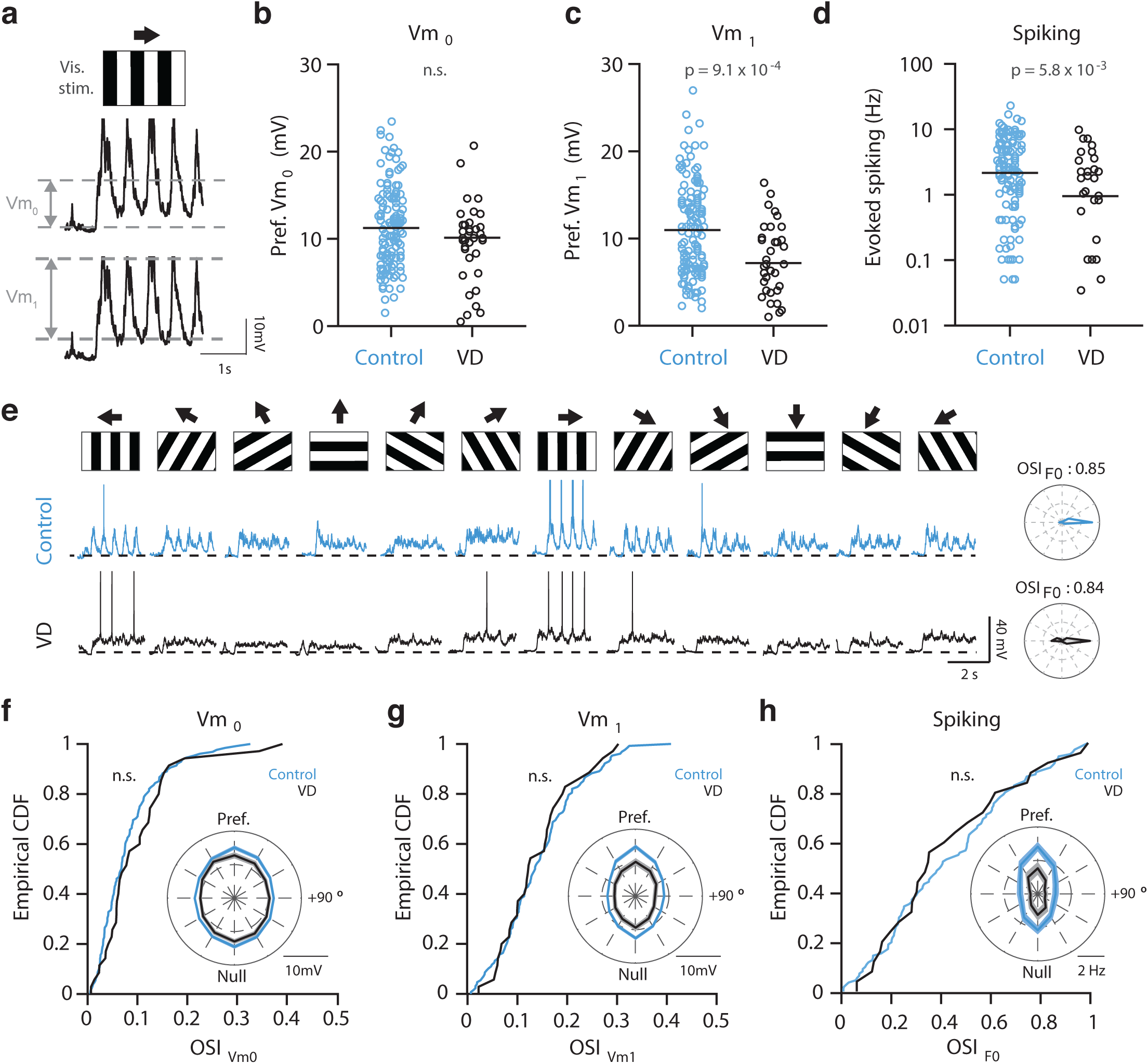
Reduced synaptic and spiking responses but no change in selectivity in visually deprived mice. (**a**) Example (spike-clipped) membrane potential trace recorded in response to a drifting grating stimulus, showing the evoked mean membrane potential depolarization (Vm_0_) and modulation (Vm_1_). Plots showing the Vm_0_ (**b**), the Vm_1_ (**c**) and spike rate in all cells that showed evoked spikes (**d**) measured at the preferred direction for spiking in control (blue) and visually deprived (black) mice. (**d**) is plotted on a logarithmic scale. (**b**-**d**) Horizontal line indicates median. (**e**) Membrane potential traces evoked by drifting gratings (moving in twelve different directions; top) for two example cells recorded in either control (blue) or deprived mice (black). Cumulative histograms showing the distributions of OSI values for Vm_0_ (**f**), Vm_1_ (**g**) and spiking (**h**). Inset: Polar plots showing the average tuning profile (aligned to the preferred direction for spiking) for all cells recorded in control (blue) or visually deprived (black) mice. Displayed is the mean and s.e.m.

Although there were significant decreases in membrane potential modulation and firing rate, cells recorded from visually deprived animals showed tuning responses that appeared very similar to those recorded in control animals (**Fig. 2e**). To examine this in detail we quantified the degree of orientation tuning of Vm_0_ (OSI_Vm0_; **Fig. 2f**), Vm_1_ (OSI_Vm1_; **Fig. 2g)** and the resultant spiking (OSI_F0_; **Fig. 2h**). Despite having been housed in complete darkness for up to five weeks, we found that the distribution of OSIs was similar between control and visually deprived mice for all OSI measures (OSI_Vm0_, p=0.26 (all neurons included); OSI_Vm1_, p=0.95 (all neurons included); OSI_F0_ p=0.96 (control: n=111, V.D.: n=25; see methods), Kolmogorov-Smirnov tests; **Fig. 2f- h**). Thus, although intrinsic and visually-evoked synaptic and spiking responses are altered, selectivity for moving gratings of different orientations remained unaffected by visual deprivation.

We next sought to identify what specific cellular parameters might contribute to this preservation of function in the absence of ongoing visual input by performing a series of correlation analyses between cellular properties and orientation tuning across the control and deprived populations. In control mice, the resting membrane potential (Spearman’s ρ=-0.51, n=111, p=1.1×10^−8^), threshold (ρ=-0.29, n=111, p=0.0018) and the distance from rest to spiking threshold (ρ=0.47, n=111, p=1.8×10^−7^) were strongly correlated with OSI_F0_ (**Fig. 3a**, **Supplementary Fig. 1a,b**). Unexpectedly, the resting membrane potential and its distance to threshold were found to be more strongly correlated with spike tuning than the degree of orientation tuning of the underlying synaptic responses (OSI_Vm0_ vs. OSI_F0_: ρ=0.35, n=111 p=1.5×10^−4^, OSI_Vm1_ vs. OSI_F0_: ρ=0.43, n=111, p=2.6×10^−6^; **Fig. 3a**). Overall, most relationships between intrinsic and synaptic parameters were similar in control and visually deprived animals (**Fig. 3a**). However, the relationship between the distance from rest to threshold and OSI_F0_ decreased significantly in the absence of visual input (V.D.: ρ=-0.03, p=0.90, n=25; vs control Fisher’s Z=2.30, p=0.021; **Fig. 3a**, **Supplementary Fig. 1a**). On the other hand, in visually deprived animals resting membrane potential maintained its strong correlation, while Vm_1_ became significantly more correlated with OSI_F0_ (control: ρ=0.30, n=111, p=0.0013; vs. V.D.: ρ=0.67, n=25, p=3.2×10^−4^; Fisher’s Z=2.16, p=0.031) (**Fig. 3a**, **Supplementary Fig. 1c**). This suggests that a cell’s resting membrane potential and membrane potential modulation are key parameters in retaining the functional profile of visual neurons (**Fig. 3b**).

**Figure 3.**
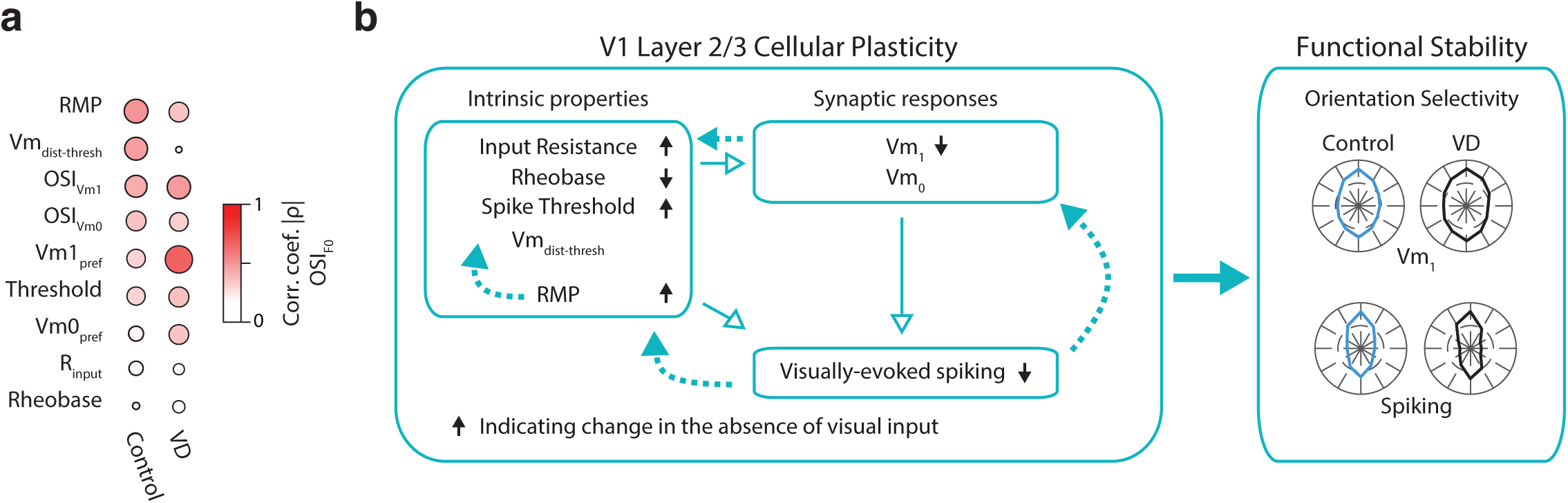
Proposed cellular mechanisms underlying the maintainance of orientation selectivity. (**a**) Plots showing the relationship between intrinsic and synaptic properties and spiking OSI in normal (left column) and visually deprived (middle column) mice. Colour and size of circle indicate the Spearman’s correlation coefficient (absolute value). Variables are ranked by correlation (p-value) in the control group. (**b**) Schematic describing the changes in intrinsic and synaptic properties in layer 2/3 neurons resulting from reduced visual input. Visual input-dependent homeostatic plasticity of these intrinsic properties (dashed arrows) is proposed to maintain robust orientation selectivity. Polar plots display an average tuning (scaled to the peak response at the preferred direction in control) obtained from all cells recorded in this study.

By performing whole cell recordings from large numbers of layer 2/3 cells in primary visual cortex we have identified several cellular parameters that are regulated by ongoing visual input. We propose that changes, such as the increase in input resistance and subsequent decrease in rheobase attempt to compensate for reduced network excitability. The cell’s resting membrane potential was identified as one key parameter that might trigger such plasticity. Various membrane potential-related sources of intracellular Ca^2+^ could drive such changes. These include spiking and backpropagating action potentials^12^ and subthreshold sources such as NMDA channels^13^ and low voltage-dependent Ca^2+^ channels^14^. Regardless of the precise mechanisms, the fact that synaptic tuning is maintained suggests such intrinsic plasticity has a global impact to preserve the relative weights of visually-evoked synaptic inputs across the dendritic tree. We propose that, under conditions of reduced sensory input, changes in intrinsic properties including resting membrane potential, trigger cellular plasticity^15,16^ to ensure functional properties such as orientation tuning are retained.

## Acknowledgements

We are grateful to the support staff of the Biological Research Facility and thank Tiago Branco and Mateo Vélez-Fort for providing comments on the manuscript. This work was funded by an Investigator Award from the Wellcome Trust (096436/B/11/Z) (T.W.M.).

## Author contributions

T.W.M. and A.P.Y.B designed while A.P.Y.B performed experiments. A.P.Y.B and L.C. analyzed the data. L.C., A.P.Y.B., and T.W.M wrote the manuscript.

---

**Figure S1.**
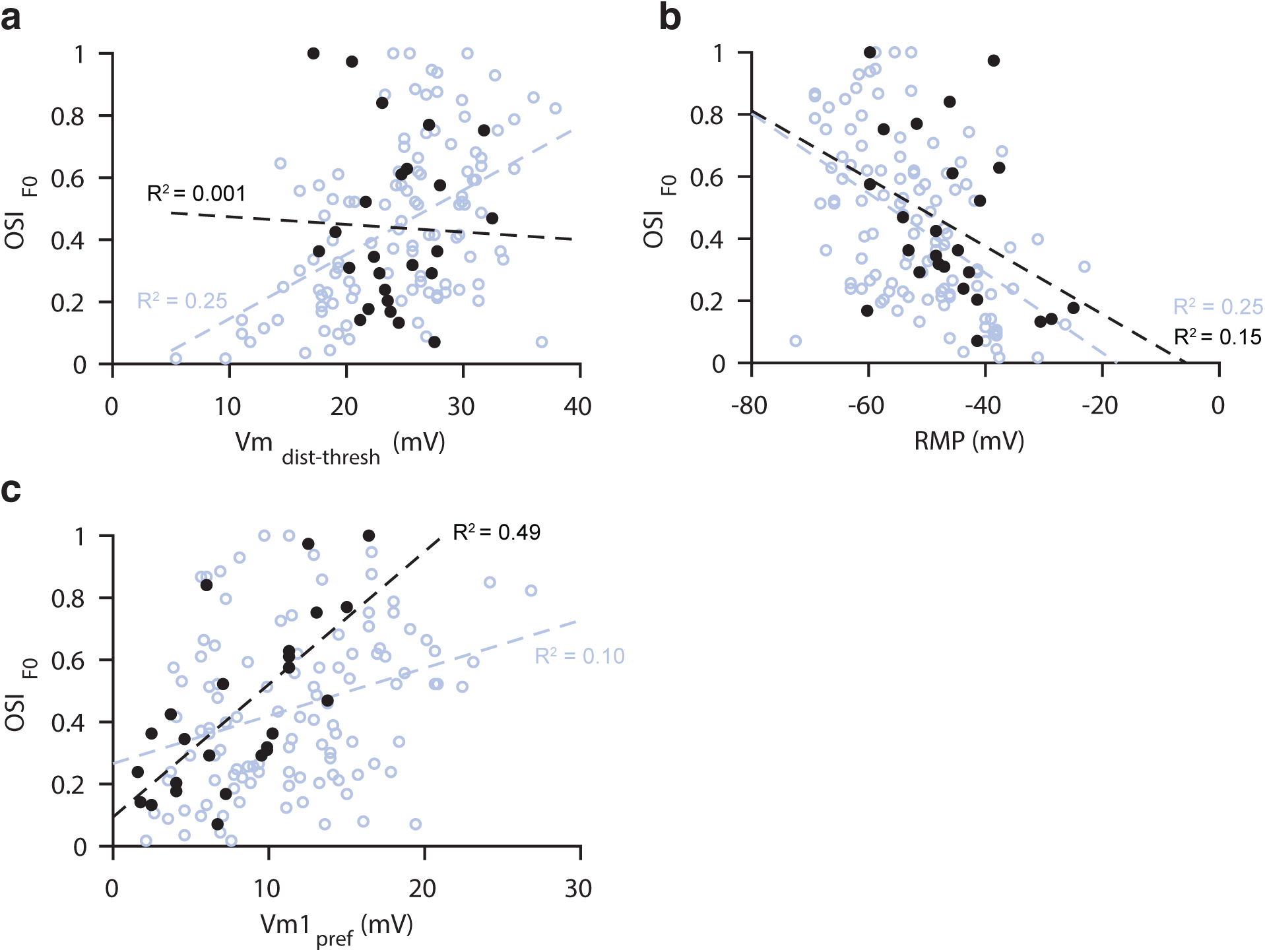
Relationship between intrinsic and synaptically evoked properties and spiking OSI. Scatter plots showing the correlations between OSI and (**a**) distance between resting membrane potential and spike threshold (Vm_dist-thresh_), (**b**) resting membrane potential (RMP), and (**c**) the membrane potential modulation at the preferred direction (Vm1_pref_). Data obtained from control (n = 111, blue) and visually deprived (n = 25, black) are overlaid with the line of best fit for each. R^2^ shown for control and deprived.

## Methods

All procedures were approved by the local ethics panel and the UK Home Office under the Animals (Scientific Procedures) Act 1986.

### Surgical procedures

Adult C57BL/6 mice (5-9 weeks old) were anaesthetized with a mixture of fentanyl (0.05mg/kg), midazolam (5.0mg/kg), and medetomidine (0.5mg/kg) in saline solution (0.9%; intraperitoneal). Anaesthesia was checked every 10-15 minutes and supplemented as necessary (20% of initial dose).

During experiments mice were maintained at 37-38°C using a rectal probe and a heating blanket (FHC, Bowdoinham, ME, USA). During surgery both eyes were protected by the application of ointment (Maxitrol, Alcon), which was then carefully removed from the right eye using a cotton bud before visual stimulation (see below). Scalp overlying the left primary visual cortex (V1) was removed and the exposed bone was carefully cleaned and roughened and then allowed to dry fully in the air. A custom-designed head fixation implant was affixed to the skull using a cyanoacrylate-based adhesive (Histoacryl, Braun) and then a dental cement mixture (Simplex Rapid, Kemdent), blackened using 2% carbon powder. During experiments the implant was fixed in a custom-made clamp.

A craniotomy, typically 1×1mm was drilled over V1 using a dental drill (Osada Electric, Japan) with a 0.3mm burr. Following removal of the bone, the exposed dura was carefully washed with cortex buffer until any bleeding had ceased. Bleeding was typically very minor; however, any blood on the surface of the brain dramatically reduced the chance of obtaining a successful gigaseal. In some animals, a small durectomy was performed before recording.

### In vivo whole-cell recordings

Patch pipettes for blind in vivo whole-cell recordings were fashioned from borosilicate glass (outer diameter: 1.5mm, inner diameter: 0.86mm, Harvard Apparatus) using a two-stage filament puller (PC10, Narashige). The resistance at the tip was 5-7MΩ; any pipettes outside this range were discarded. The tip size of such pipettes was approximately 1.5μm.

Intracellular solutions for whole-cell recordings was prepared in batches of 2x concentration and frozen in single-use aliquots at −20°C for one month or less, or - 80°C for longer periods. The final concentrations (1x) were (all from Sigma-Aldrich or VWR International, UK; in mM) 110 K+, 8.5 Na^+^, 5 Mg^2+^, 0.04 Ca^2+^, 110 MeSO_3_^−^, 12.04 Cl^−^, 0.05 EGTA, 40 HEPES, 4 ATP, and 0.5 GTP; the pH was adjusted to 7.28 using KOH and/or HCl. On the day of the experiment, the stock was diluted to 1x, and measured for osmolality (mean 289mOsm±3.64, range 281-295mOsm) using a vapour osmometer (Vapro 5520, Wecor). Once prepared the solution was kept on ice for the duration of the experiment.

Pipettes were placed in a holder attached to a preamplifier (HS-2A, Axon), mounted on a manipulator (4-axis Junior, Luigs & Neumann) and connected to an Axoclamp 2B amplifier (Axon). Moderate pressure was applied as pipettes were lowered to the brain surface under a 10x water-dipping objective (Olympus), and then rapidly advanced to ∼200μm from the pial surface. Positive pressure was then reduced to approximately 30mbar and cell search carried out in voltage-clamp mode (Margrie et al. 2002).

Once whole-cell access was obtained, the amplifier was switched to current-clamp mode and the series resistance was compensated for manually using a bridge circuit. Access resistance was in the range of 15-55MΩ (mean 30.3±9.4MΩ, n=150). All the experiments described here were carried out in current-clamp mode, with no holding current. Junction potential was not corrected for. Data were low-pass filtered by the amplifier at 10kHz and acquired at 25kHz using an ITC-18 interface (Instrutech) using IGOR Pro software (Wavemetrics) running the Neuromatic package (available at http://www.neuromatic.thinkrandom.com/). Electrical (50Hz) noise was minimised by passive electromagnetic shielding of the sample and pre-amplifier. Additionally, a HumBug device (Quest Scientific) was used to further reduce any residual 50Hz electrical noise.

### Response to current injection

Once stable access to a cell was obtained and the bridge compensation set, an IV protocol was carried out. Square wave current pulses of 1000ms duration were applied, beginning with a hyperpolarising step of −400pA with decrements of 50pA, until the 9^th^ step in which no current was injected. From the 10^th^ step, depolarising current steps with increments of 25 pA were used. Supra-threshold current steps were carried out up to at least 1.5x rheobase.

### Visual stimulation

Visual stimulation was carried out using an 8” monitor (ADP-1081AT, DataSound Laboratories) positioned 9cm from the animals left eye, subtending approximately ±42° in azimuth and ±34° in elevation, positioned at ∼45° to the long body-axis. A photodiode was positioned over the lower-left corner of the monitor and data were acquired in parallel with electrophysiological recordings to provide an accurate stimulus timestamp. Stimuli were generated using scripts written in MATLAB (Mathworks) using the Psychophysics Toolbox (version 3, http://psychtoolbox.org/). Stimuli were generated on a dedicated computer to ensure reliable performance.

Drifting grating stimuli consisted of square wave full-contrast gratings. Grating stimuli consisted of a stationary oriented grating (‘hold’) for 2s, followed by a sustained period of drifting grating (2.5s), followed directly by the next stationary grating. 12 evenly spaced directions were presented with a spatial frequency of 0.0283 cycles per visual degree, and a temporal frequency of 2 cycles per second. Between three and 15 repetitions of the 12 directions were presented.

### Visual deprivation

For experiments involving visual deprivation, animals were transferred to cages with fully blackened walls at post-natal day 19 (P19), immediately after weaning (but after eye-opening which occurs at P13-14) and thus includes the critical period (Levelt & Hubener, 2012). The cages were blackened with several layers of matt black emulsion, and covered with a blackout curtain (Thorlabs) to ensure no light could enter around the edges. The blackening of the cages was tested using a light meter (X-Cite XR2100), revealing that minimal light was transmitted through the cage walls even in strong sunlight (<0.005 µW/m^2^). Animals were kept in full darkness until the experiment day, with the exception of brief checks for food and water replenishment which were carried out under low intensity red light.

On the day of the experiment, animals were anaesthetised under red light and placed in a black-out box until the anaesthetic had taken full effect. The eyes were then covered with cream then were further covered in with blackout material to avoid exposure to surgical lighting. This material was not removed until immediately prior to the experiment.

Experiments on visually deprived (VD) animals were performed after at least 18 days of visual deprivation (P37), and up to P53. As the age profile of the VD animal cohort did not precisely match that of the control dataset, all variables described were tested for correlation with age; none were significantly correlated.

### Data Analysis

All experiments were logged in a central database (FileMaker Pro, FileMaker) and were analyzed using MATLAB (Mathworks). Electrophysiological data were imported from Igor PXP format and read in to MATLAB using a custom-written package, based on a similar R package (available at: https://github.com/jefferis/IgorR). Action potentials (APs, ‘spikes’) were detected using a two-step algorithm. First, spikes were detected by finding peaks of dV/dt greater than a threshold of 8x the standard deviation of a 100ms baseline period at the start of the trace. Spike threshold was defined as the membrane potential (Vm) at maximal d^2^V/dt^2^ up to 4ms before this point. Results of the spike detection procedure were manually verified in all cases. Before any analysis of intrinsic or evoked membrane potential properties, spikes were clipped by removing data points from 1ms before spike threshold till 10ms after and linearly interpolating. All references to Vm values/changes in this text refer to spike-clipped data.

In order to classify regular spiking neurons, we analyzed the waveform of the first evoked AP at rheobase. The only parameters used in classification were the amplitude and time-to-peak of the after-hyperpolarisation (AHP). AHP amplitude was defined as the peak hyperpolarisation within 10ms of the peak of the spike, relative to the steady-state membrane potential (Vm_ss_) of the neuron at rheobase. Vm_ss_ (the mean Vm between 250 to 50ms before the end of the IV current-step) was used as the baseline measure for calculating AHP amplitude rather than spike threshold, as some neurons displayed a prominent, long-lasting ramp potential before reaching spike threshold, which then artificially increased the estimate of AHP amplitude.

Neurons were classed as regular spiking (RS) if their AHP amplitude was <5mV, and occurred within 10ms following the peak spike depolarization. All the data in this study were therefore obtained from regular-spiking neurons.

For analysis of intrinsic properties (Figure 1) we performed the following analyses. The resting membrane potential (RMP) was taken as the Vm_ss_ of the voltage trace when no current was injected. Rheobase was taken as the first positive current step to elicit a spike. The spike threshold was taken as the threshold of the first spike observed at rheobase. Hyperpolarizing input resistance was calculated at −200pA current injection step. Depolarizing input resistance was calculated at 150pA current injection step and only neurons with a rheobase >150pA were included in the calculation (control: n=128, V.D: n=35 cells).

For analysis of visually-evoked responses (Figure 2), the membrane potential response to each direction was divided into a “baseline” window (the final 500ms of the static grating stimulus, before stimulus drift onset) and a “stimulus-evoked” window (0.5-2.5s after drift onset).

The mean membrane potential response of a neuron to stimulation (*Vm* _0_) was defined as follows. For each trial, *i*, for each direction, *θ*, (*Vm*_0_) *i*,*θ* was calculated as:

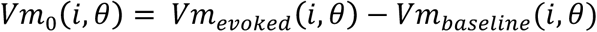

where *Vm*_*baseline*_ is the mean membrane potential during the baseline window, and *Vm*_*evoked*_ is the mean membrane potential during the stimulus-evoked window.

One concern with this definition is that *Vm*_*baseline*_ may differ in an orientation selective manner - in other words, that a neuron’s steady-state membrane potential is tuned for stationary grating stimuli. Such an orientation-dependent baseline measure may introduce an apparent tuning in a cell not otherwise tuned for drift orientation, or mask the orientation tuning of a cell which is tuned. To investigate this possibility, the probability of the baseline of each neuron being tuned was calculated using a one-way ANOVA, testing the hypothesis that the baseline differed across orientation conditions. At the 0.05 significance level only 3/128 were found to have a baseline which varied across conditions, fewer than the predicted number of type I errors. Indeed, fewer cells were found to have a stimulus-responsive baseline than would be predicted by the type I error rate at all significance levels. Therefore, it is unlikely the baseline measure used here influences the tuning for orientation as measured to drifting gratings.

The membrane potential modulation of a neuron (*Vm*_1_) was quantified as follows. For each direction, the membrane potential responses during the stimulus-evoked window was averaged across trials. Next, the stimulus-evoked window was divided into four separate regions (each 0.5s long) and the traces in these regions were averaged (giving the average response to a full cycle of the drifting grating). Finally, the membrane potential modulation (*Vm*_1_) was taken to be the voltage difference between the maximum and minimum of the averaged trace.

Neurons were classified as responsive to drifting gratings based upon whether changes in mean firing rate during the drifting gratings were statistically significant. First, spiking responses, *F*_*0*_, were calculated, for each trial, i, and each direction, *θ*, as:

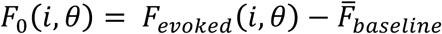

where 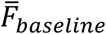 is the firing rate across all baseline periods (all trials, all directions), and F_*evoked*_ is the firing rate during the stimulus-evoked window (thus, *F*_*0*_ can take negative values). Next, for each neuron, a Wilcoxon sign-rank test was performed on the distribution of |*F*_*0*_| values, to determine whether the stimulus-evoked firing rate was significantly different from baseline.

To calculate the Orientation Selectivity Index (OSI) we used vector methods. First, we computed the normalized vector average of the responses over orientation space (Swindale, 1998; Ringach et al. 2002):

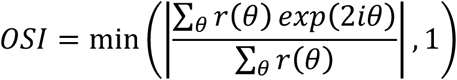

where r(**θ**) are responses to each direction, *θ*. Three measures of OSI were calculated. OSI_F0_ was calculated by taking *r*(*θ*) to be the trial-averaged spiking responses (*F*_*0*_):

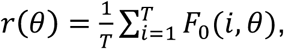

OSI_Vm0_ was calculated by taking the trial-averaged mean membrane potential response (*Vm*_0_):

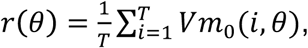

and OSI_Vm1_ by taking *r*(*θ*) to be the membrane potential modulation (*Vm*_1_): as computed above:

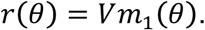

Only cells classified as responsive to drifting gratings (see above) were used to calculate OSI_F0_ (control: n=111, V.D.: n=25). The OSI could exceed 1 (due to negative responses after baseline subtraction) and in such cases we set the value to 1 (control: 3/111 cells, V.D.: 0/25 cells).

In order to consider responses at the preferred direction, preferred direction was calculated in the following way. The preferred orientation was taken as:

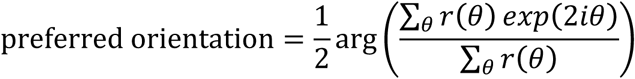

Responses were then considered at the grating directions closest to the preferred orientation and to the preferred orientation+180°. The preferred direction was defined as the grating direction that elicited the largest response. For example, a neuron with a preferred orientation of 20° would have a preferred direction of 210° if the response to drifting gratings at 210° exceeded the response to gratings at 30°; otherwise the preferred direction would be defined as 30°.

### Statistical Analysis

All statistical analyses were carried out in MATLAB. The median and interquartile range are reported when comparing the intrinsic and evoked properties of control and visually deprived groups. In order to compare these medians statistically we used the Wilcoxon rank-sum test. For all correlation analyses we used Spearman’s rank-order correlation. In order to test for differences in correlation values between control and visually deprived groups we used a direct application of Fisher’s Z-transformation. In order to test for differences in the OSI distributions between control and visually deprived groups we used the Kolmogorov-Smirnov test.

